# Estimating the abundance of duiker (*Cephalophus spp.*) populations in African rainforests: an overview of relevant methods

**DOI:** 10.1101/556332

**Authors:** Gaius Bolumbu Entanga Elenga, Jean-Michel Gaillard, Dieudonné Eyul’ Anki Musibono, Séraphin Ndey Bibula Ifuta, Christophe Bonenfant

## Abstract

Duikers are among the most sought after antelope species for bush meat in central Africa. Estimates of population abundance of duikers based on reliable methods is therefore of prime importance for their sustainable management. Here we retrieved 31 studies from the literature and compared methods used to estimate the abundance of duiker populations in African rainforests. Implemented methods all derived from seven main combinations of sampling designs and population abundance estimators. We then evaluated the relevance of those seven methods by scoring them based on eight criteria selected according to their pros and cons reported in the litterature for large-scale population management of wildlife. For management purposes, methods derived from distance sampling offer the best compromise between the implementation costs and the biological information collected. In particular, both diurnal and nocturnal distance sampling can be recommended. Hunter calls and drive-netting are less reliable, but can be used in association with other measurements in the framework of indicators of ecological changes, a monitoring approach that has been successfully used in temperate ecosystems for managing large herbivores.

**Funding information:** Ministère de l’Environnement et du Développement Durable de la République Démocratique du Congo, Campus France and Centre National de la Recherche Scientifique (CNRS).

**Résumé:** Les céphalophes sont parmi les espèces d’antilopes les plus recherchées pour la viande de brousse en Afrique centrale. L’estimation de l’abondance des populations de céphalophes revêt donc une importance primordiale pour leur gestion durable et adaptative. Nous avons extrait 31 études de la littérature et comparé les méthodes utilisées pour estimer l’abondance des populations de céphalophes dans les forêts tropicales africaines. Les méthodes employées dérivent de sept combinaisons principales de plans d’échantillonnage et d’estimateurs d’abondance. Nous avons évalué la pertinence de ces sept méthodes sur la base de critères sélectionnés en fonction de leurs avantages et inconvénients pour une gestion à grande échelle de la population. Les méthodes dérivées de l’échantillonnage par distance offrent le meilleur compromis entre les coûts de mise en œuvre et les informations biologiques collectées. Les méthodes d’échantillonnage diurne et nocturne se sont montrées les plus satisfaisantes. Les méthodes d’appel par les chasseurs et de battue au filet sont moins fiables, mais peuvent être utilisées en association avec d’autres mesures biologiques comme indicateurs de changements écologiques, une approche qui a été utilisée avec succès dans les écosystèmes tempérés pour la gestion des grands herbivores.

**Mots-clés:** gestion durable, grands herbivores, indicateurs de changement écologique, méthode de recensement, viande de brousse

## 1 INTRODUCTION

Over the last centuries, human activities have progressively became the most influential driver of biodiversity dynamics (Dirzo et al., 2014) and among the strongest evolutionary force of living organisms (Palumbi, 2001). Many wild animal populations of vertebrates are exploited either for sport (Gordon, Hetser & Festa-Bianchet, 2004) or for subsistence (Milner-Gulland & Resit-Akçakaya, 2001). Mirroring the human population growth, the African fauna is the target of an intense and widespread exploitation by humans outside of protected areas (Lahm, 1993; Nasi & van Vliet, 2011). Trophy hunting for the big five is still the aim to achieve for wealthy hunters traveling to the African continent (Di Minin, Fraser, Slotow & MacMillan, 2012). Besides, most mammals and particularly large herbivores have been part of the human diet since ever (Caro & Riggio, 2014), and this practice is still persisting nowadays (Ripple et al., 2016). For instance, Nasi, Taber & van Vliet (2011) estimated to 4,5 million metric tons of wild meat per year the biomass harvested in the rainforests of the Congo basin, central Africa. The estimated volume of bush meat consumed in this area ranges between 1 and 5 million metric tons per year (Fa & Brown, 2009; van Vliet et al., 2012; Wilkie & Carpenter, 1999). Overall, bush meat makes a substantial share of animal protein sources in many African countries (Rowcliffe, 2002; Fargeot, Drouet-Hoguet & Le Bel, 2017) and this often illegal activity contributes to the economy of both rural and urban human populations (Lescuyer & Nasi, 2016).

For most activities related to bush meat production including hunting, marketing and consumption, duikers (*Cephalophus spp.* Hamilton Smith, 1827) usually rank first in terms of relative abundance in comparison with other species of large herbivores. Consequently all species of duikers are under great anthropogenic pressure, being much consumed and heavily traded by indigenous people and local communities, although rural and urban populations are no exception to this general picture (Bahuchet & Loveva, 1999; Fargeot, 2006; Newing, 2001; Semeki et al., 2014; van Vliet & Nasi, 2008). Currently, duikers belong to either the LC “minor concern” category or the NT “near-threatened species” category, according to the IUCN classification (CITES, 2018; UICN, 2019). However if duikers are currently not critically endangered, they could rapidly become so if their trading and exploitation are not under control and their populations not managed properly in the wild. For instance, past studies suggest that the reduction of duiker density in forested areas where hunting is practiced compared to no-hunting areas could reach 12.9% – 42.1 % in Ituri, Democratic Republic of Congo (Hart, 2000), 43 – 100% in Makokou, Gabon (Lahm, 1993; van Vliet & Nasi, 2008) and 43.9 % in Massapoula in the Central African Republic (Noss, 1999).

Duikers *sensu largo* are small- to medium-sized ungulates mainly living in African forests. Those species are good indicators of habitat modification and some of them are ‘pioneer’ species *i.e.* species rapidly colonizing habitats at early stages of the vegetation succession (Dubost, 1980; Newing, Davies & Linkei, 2004). On the African continent duikers correspond to at least 16 known species (Wilson & Mittermeier, 2011) that represent one of the main sources of animal protein for human populations inhabiting forested areas. Being part of the ‘common biodiversity’ (*sensu* Hanski, 2005), duikers attract little attention to wildlife managers and conservation agencies. For instance, estimates of duiker population growth rates, which could be used as a starting point for management, are scarce in the literature. Overall, the ecology of duikers remains surprisingly poorly known and particularly so their population dynamics. From an ecological point of view, duikers constitute an important part of primary consumers of the rainforests. By consuming tree seeds (*e.g. Irvingia gabonensis, Pseudospondias longifolia, Dacryodes buettneri, Dacryodes edulis, Ancystrophyllum spp.*), they play a key role in seed dispersal and tree dissemination (Dubost, 1980; Gautier-Hion, Emmons & Dubost, 1980; Feer, 1989, 1995). In addition, duikers are important prey species for some carnivores such as leopard (*Panthera pardus*), African golden cat (*Caracal aurata*), African civet (*Civettictis civetta*) or African wild dog (*Lycaon pictus*).

In addition to a lack of biological knowledge on duikers and to the extensive pressure humans put on their populations, many socio-economic factors make duiker management difficult in practice. Among the myriad of problems that slow or even prevent the implementation of sustainable hunting in the different tropical forest ecosystems of the Congo Basin in central Africa, six are particularly acute and likely to have a strong impact on the future prospect of duikers: obsolete management rules and policies (van Vliet & Nasi, 2017), illegal logging and mining (Nasi et al., 2008), high human population growth (van Vliet et al., 2012), non-compliance with hunting legislation (Fargeot, 2006), poverty of the population (Kumpel, 2006; Lescuyer & Nasi, 2016) and climate change (Nasi et al., 2008). As the most visible ecological consequence of this unmanaged exploitation, African forests are progressively emptying of their duikers, which display a severe contraction of their original distribution over the last decades (Nasi, Taber & van Vliet, 2011). For management purposes but also for conservation we are facing an urgent need and demand to propose objective and reliable tools for monitoring duiker populations and assessing their current status.

To this end, several methods have been proposed and used to estimate the abundance of duiker populations in rainforests (Table 1; Fig. 1). In the absence of common questions and management goals, methods previously employed to assess population abundance have varied a lot in terms of sampling designs and ability to account for imperfect detection of animals (Seber, 1982; Nichols 1992; Schwarz & Seber, 1999). For instance, White & Edwards (2001) carried out sampling of duikers based on direct observations while Omasombo, Bokela & Dupain (2005) relied on indirect measures of occurrence and number of feces. However, regardless of the methods used the reliability of the population abundance estimates derived from these methods is questionable (Koster & Hart, 1998; Foster & Hansen, 2012; Mathot & Doucet, 2006; Walter et al., 2006). Indeed, the secretive behaviour and nocturnal habits of some species such as the Bay duiker (*C. dorsalis*), the poor visibility conditions within habitats used by duikers (canopy openness, presence of understory and of tall grass), the strong spatial variation in human disturbances (*e.g.* hunted vs non-hunted areas, local deforestation) all affect the detection rate of duikers (Fa & Brown, 2009; van Vliet & Nasi, 2007). Moreover the relevance of those seven different sampling designs and methods used to estimate duiker abundance in a population management context has not been assessed so far.

**TABLE 1:** Authors and sampling methods used for estimating abundance of duikers in central African rainforests.

| **Authors** | **Sampling Method** |
| --- | --- |
| van Vliet et al., 2009; Wilson, 1990. | Hunter call method |
| Dubost, 1980; Newing, 1994; Noss, 1999. | Drive-netting method |
| Elenga et al., 2013; White, & Edwards, 2000; Kamgaing et al., 2018; Lahm, 1993; Payne, 1992; Walter et al., 2006. | Nocturnal distance sampling |
| Koster, & Hart, 1988; Lannoy et al., 2003; Plumptre, 2000; Griffiths, 1978; Jost Robinson, Zollner, & Kpanou, 2016. | Diurnal distance sampling |
| Mathot, & Doucet, 2006; van Vliet et al., 2009; Wilkie, & Finn, 1990; Rovero, & Marshall, 2004. | Dung count method |
| Forboseh et al., 2007; Omasombo, Bokelo, & Dupan, 2005; Maréchal, & Bastin, 2008; Tutin, White., & Mackanga-Missandzou, 1997; Walsh & White, 1999. | Recces, recognition walk or sweep method |
| Bowkett et al., 2007; Carbone et al., 2001; Foster, & Harmsen, 2011; Troillet et al., 2014; Rovero, Jones, & Sanderson, 2005. | Camera trap method |
| Prins, & Reitsma, 1989. | Tracks count method |

**FIGURE 1:**
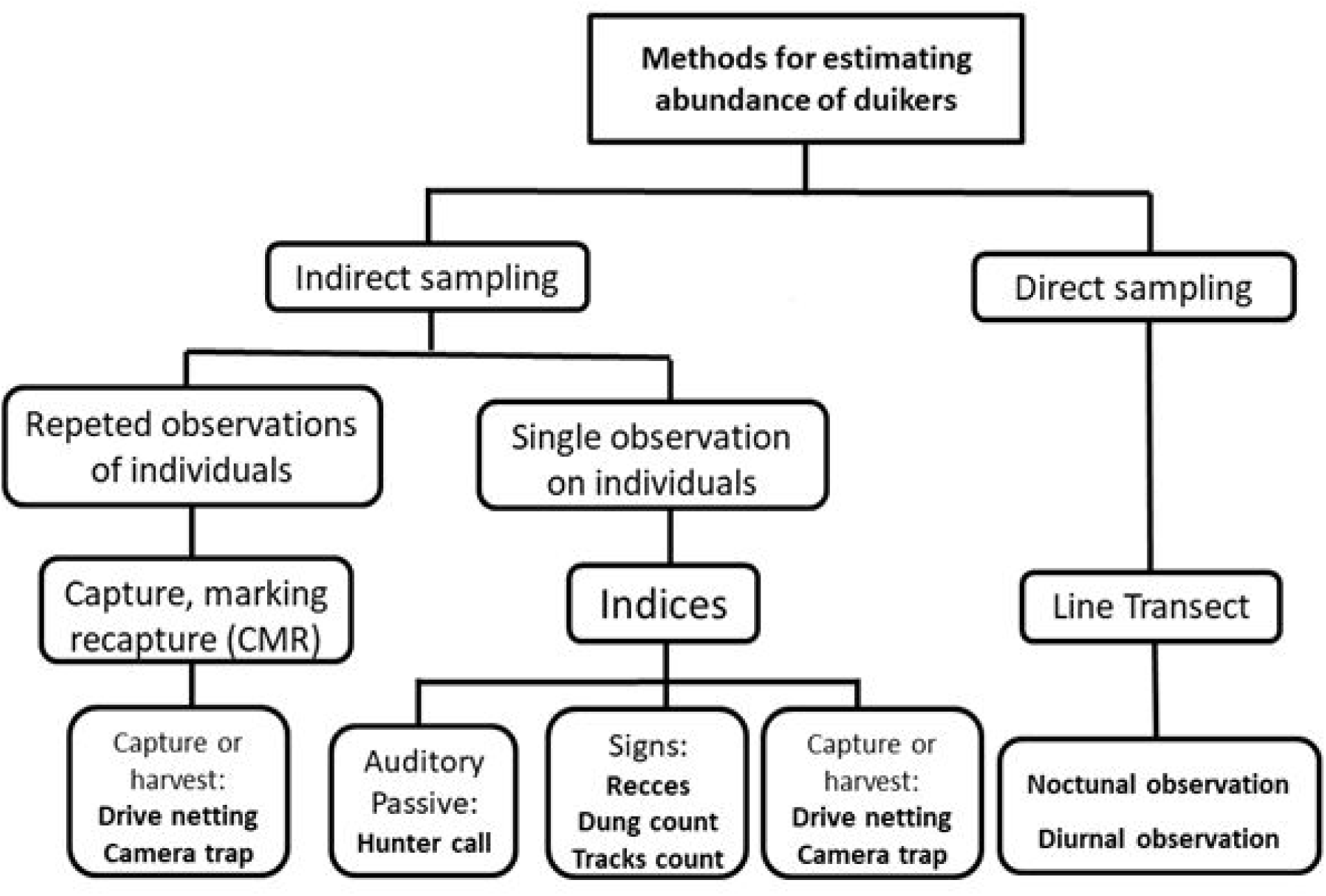
Overview of the different methods and sampling techniques used to estimate abundance of duikers in central African rainforests. A variety of approaches have been implemented in the field using either direct or indirect sampling methods such as hunter call. Most importantly, many methods provide relative abundance estimates only such as the indirect sampling methods by single observation on individuals based the indices (i.e. hunter calls and drive-netting methods), while few others did indeed returned an absolute estimate of duiker abundance by estimating detection rate of duikers.

In this study, we aim to fill the gap by first providing a review of the published studies on the different methods and sampling designs that have been used to estimate duiker abundance. We then assess the relevance of these methods to identify which methods offer the best compromise between cost-efficiency and reliability to estimate duiker population abundance given the difficult observation conditions of the African rainforest. We lastly discuss how some simple, easy-to-do methods may be combined to additional biological measurements collected on harvested duikers such as body mass or jaw length to propose an adaptive management tool for duiker populations.

## 2 MATERIAL AND METHODS

### 2.1 Literature search

We searched the literature from three different sources of information: grey literature, reference books and online consultation of databases and catalogs. We retrieved published material from four online search engines: (*i*) Google scholar for its flexibility and ability to recover reports and grey literature; (*ii*) the online database of the Congo Basin Forest Partnership (CBFP: http://www.pfbc-cbfp.org/actualites/items/EDF_2010-F.html). This latter site has provided references based on news (recent and past) from the Congo Basin countries in Central Africa; (*iii*) the use of the APES online database "Ape Populations, Environments, and Surveys" and "IUCN-New Population, Environments and Surveys Database" (http://apes.eva.mpg.de/ and http://www/iucn/org/en/news_homepage/news_par_date/3246/New-ape-population-environments-and-surveys databases). This site has provided, apart from the results on primates, several reports of inventories of large mammals made in the forest; and finally, (*iv*) the use of international publication databases including Ingenta Connect, Wiley InterScience, SpringerLink and Unicat. We queried all databases and catalogs with the following combination of keywords: “census” “methods”, “monitoring”, “survey”, “Congo basin”, “mammal”, “herbivorous mammals”, “ungulates”, “fauna", "density-ies", "duiker" back to 1970.

### 2.2 Assessing the relevance of the methods used to estimate abundance

The approach used was inspired and adapted from several authors (Jachmann, 1991; Jachmann, 2002; Thorn et al., 2010; Whitehouse, Hall Martin & Knight, 2001). As a first step, we defined eight criteria to classify the methods used to estimate duiker abundance for management purposes: (c1) the representative sampling design; (c2) the ability of the method to identify the duiker species; (c3) the influence of environmental factors on animal detection (*i.e.* food availability, local microclimate, undergrowth density, hunting pressure); (c4) the risk of double counts; (c5) the level of technicality to be deployed in the field; (c6) the time required for data collection; (c7) the money costs of the method in terms of logistic and manpower; (c8) the ability of the method to produce an estimate of population density with an associated measurement error.

In a second step, we scored the methods based on the chosen criterion weighted according to its degree of relevance for management or research. The weight assigned to each criteria varied from 0 to 3 as follows: 3, if the criterion was "very important"; 2, if the criterion was "important"; 1 if the criterion was “less important” and 0, if the criterion was “not important” (Table 2). The product of the criteria weight and associated scores returned the final score of the different abundance estimation methods. The final score was calculated so that the greater it was, the most suitable and relevant it was for duiker population management.

**TABLE 2:** Criterions used to evaluate methods of abundance estimation for management of according of duikers in African rainforest. The different pros and cons we extracted and summarized from a literature survey. To compute an overall score of each method of abundance estimation, we weighted each criterion from 0 to 3 depending on it relevance for wildlife management. For instance, a cheap method is favored over an expensive one so that the costs of implementation is given a weight of 3 (criteria c7).

| Criterion (c) | Unweighted | Weighted to management |
| --- | --- | --- |
| c1, the method have a good sampling design.<br>We suppose that all methods are good sampling<br>(c1 = 1) | 1 | 1 |
| c2, the ability of the method to identify duiker:<br>very difficult ( = 0), difficult ( = 1), good ( = 2)<br>and very good ( = 3) | 1 | 3 |
| c3, the influence of the method on<br>environmental factors (low visibility, local<br>microclimate, dry season, undergrowth density, ,<br>rainy season, strong wind,etc.): very sensitive ( = 0),<br>sensitive ( = 1), less sensitive ( = 2) and<br>not sensitive ( = 3) | 1 | 2 |
| c4, the risk of method counting twice the same<br>species: very high ( = 0), high ( = 1), lower ( = 2)<br>and almost zero ( = 3) | 1 | 3 |
| c5, level of technicity: complex ( = 0), difficult ( = 1),<br>easy( = 2) and very easy ( = 3) | 1 | 3 |
| c6, the time used by the method to collect data<br>and information, including analysis and<br>processing; very long ( = 0), long ( = 1), short ( = 2)<br>and fast ( = 3) | 1 | 2 |
| c7, the material and financial resources required by the method: very expensive ( = 0), expensive ( = 1), less expensive (= 2) and not expensive ( = 3) | 1 | 3 |
| c8, the ability of the method to estimate abundance populations combined with a measurement error: very good ( = 3), good ( = 2), less good ( = 1), and almost zero or very bad ( = 0) | 1 | 1 |

Key: « 0 »: not important; « 1 »: less important; « 2 »: important; « 3 » very important;

## 3 RESULTS

We recovered 31 publications that addressed the question of duiker abundance estimation methods. We identified seven combinations of sampling designs and methods to estimate duiker abundance: (1) Hunter call; (2) camera trap; (3) animal drive or drive-netting; (4) nocturnal distance sampling; (5) diurnal distance sampling; (6) dung counts; (7) recognition walk or indirect *Recces;* and (8) tracks count. These methods can be divided into two broad categories (Fig. 1) according to Eberhardt (1978): direct sampling vs. indirect sampling depending on whether individuals are mobile and detected with certainty, or not. We briefly describe below these different methods.

The distance sampling (Buckland et al., 2001) used in methods 4 to 6 consists in walking parallel tracks in the forest in the same direction, then collecting data along these transects. The observations collected are the distance perpendicular to the transect of each seen duiker. Distance sampling is the only direct sampling methods suitable for moving animals (Eberhardt, 1978) and has been applied to duikers using two different sampling designs: (*i*) diurnal distance sampling method (DSM) when the observations are direct and take place during the day, the morning between dawn and 11:00 am or between 03:00 pm and 06:00 pm (Jost Robinson, Zollner, & Kpanou, 2016; Koster et al., 1988; Lannoy et al., 2003; Newing et al., 2004); (*ii*) Nocturnal distance sampling method (NSM) when the observations are direct and take place at night, from 08:00 pm to 04:00 am the following day, a time period when animals are most active. Duikers are detected by walking slowly in the forest with headlights, searching for light reflection by the eyes of these nocturnal animals. Duikers usually freeze and the tracker approaches to identify species (Elenga et al., 2013; Kamgaing et al., 2019; Payne, 1992; Walter et al., 2006); (*iii*) Distance sampling of presence indices (DSI) when observations are indirect and based on the sign of animal presence such as dung, hair, tracks or footprints (Koster & Hart, 1988; Lannoy et al., 2003; Plumptre, 2000; Prins & Reitsma, 1989; Rovero & Marshall, 2004). Although some authors suggested this could lead to estimate absolute abundance of duikers, we classified this method among those measuring relative abundance.

Most of the time, however, the authors relied on an indirect sampling because duikers, like most mobile animals, cannot be enumerated. Among those, capture-mark-recapture (CMR) methods and derivatives returns an absolute abundance estimation (Schwarz & Seber, 1999). CMR methods rely on the identification of animals (by the mean of natural or artificial marks) from which a recapture rate can be estimated and used to estimate population size (Nichols, 1992). We found one study based on CMR, namely the drive-netting method (DNM), involving physical capture and handling of animals achieved by isolating a large area with nets. Once set, hunters enter in the area and drive the animals towards the nets. Captured duikers are identified with individual marks, and then released and eventually captured or seen again later on (Noss, 1999; Newing et al., 2004). To mark duikers, fluorescent dyes or marks on the coat, horns, mane, or legs have been used. Although most of these marks share the pitfall of being labile in time they are sometimes the only way to identify directly duikers in dense forest (Lannoy et al., 2003; Payne, 1992; White & Edwards, 2001).

Besides distance sampling and CMR methods, we retrieved many different sampling designs providing managers with relative abundance methods. Most of these methods make the assumption of a constant and proportional number of animals seen or signs of presence and the true, unknown abundance (Nichols, 2002; Anderson, 2001). Because the detection rate of individuals or signs is not estimated, these methods cannot assess population size or density of duikers but only return a frequency of observations per area or time unit. In the present review, we identified five different methods to estimate relative abundance of duikers. (*i*) The hunter call method (HCM) consists in producing an attractive sound for duikers that lasts approximately two minutes, repeated on successive determined positions. After a few minutes, because of their territorial behaviour, duikers respond to the call by moving toward the person who produced the call. Abundance is generally expressed in terms of success rate (%) to duiker calls (van Vliet et al, 2009; Newing, 1994; Wilson, 1990). (*ii*) Indirect recces method (IRM, also referred as REconnaissanCE or "ikis" or "eccel”) consists in recording the indirect observations of the animals (feces, dung, animal footprints, etc.) by walking pre-existing tracks of least resistance in the landscape, either in a pre-defined direction ("guided" reconnaissance) or without precise heading (Recognition "trip") (Forboseh, Sunderland & Eno Nku, 2007; Maréchal & Bastin., 2008; Walsh & White, 1999). The distance perpendicular to observations are not measured and the sampling plan is not important (Newing et al., 2004; White & Edwards, 2001), so the method returns a frequency of signs per distance unit. (*iii*) Camera trap method (CTM) or photographic trapping method relies on the observation of duikers from photographies. Camera traps are set in places supposedly frequented by animals (Bowkett, Rovero & Marshall, 2007; Carbone, Rovero & Marshal, 2001; Foster & Harmsen, 2012; Rovero, Jones, & Sanderson, 2005; Troillet et al., 2014) or systematically according to a geographic grid, so as to offer systematic coverage of the entire study area and allow simultaneous monitoring of multiple species (Swanson et al., 2015). Unless additional assumptions about animal movement are met, CTM yields a number of animals per time unit. (*iv*) The dung count method (DCM) records indirect signs of animal presence along linear transects (*e.g.* Putman, 1984). Again, this method gives a number of indices per distance unit and hence corresponds to a measure of relative abundance. (v) The Tracks count method (TCM) records footprints left on the ground by animals either on the road or in forest transects. Assuming a guess estimates of the duration of tracks in time, some authors derived crude estimates of densities of duikers (Prins & Reitsma, 1989). Although it does not solve the imperfect detection issue, we noted a few attempts to combine different approaches such as Recce and transects (the so called sweep sampling methods) to increase the speed of progress and to calibrate the results (Tutin et al., 1997; Maréchal, 2011) but essentially returns a number of duikers seen per surface unit instead of distance unit.

On the basis of our eigh criteria, we found that distance sampling methods offered the best compromise between costs of implementation and collected biological information for estimating abundance of duikers in the tropical rain forests (Table 3). The nocturnal distance sampling (NSM) ranked slightly higher than the diurnal distance sampling (DSM) with scores of 66.6% vs. 58.3%, respectively. The hunter call method (57.0%) and drive-netting (54.1%) ranked a bit lower. At the other end, dung counts (DCM) and tracks count (TCM) on transects are not to be recommended for wildlife management with a final score of 34.7% vs 36.1% respectively. A principal component analysis (PCA) performed on the criteria scores of all methods accounted for 83.4% of the total inertia (first 2 principal components) and highlighted how the different criteria covaried among methods used to assess duiker abundance (Fig. 2). The first principal component (PC1) was associated to criteria c8, c4 and c2 and opposed the distance sampling and drive-netting methods estimating absolute abundance, which are also costly and of high technicality, to simpler methods estimating relative abundance. The second principal component (PC2), associated to criteria c3, c5, c6 and c7, ranked methods along a continuum going from cheap and quick to carry methods to more time-consuming ones.

**TABLE 3:** Scores of survey methods for estimating duiker populations in central African rainforest. We scored the methods by weighted the 8 criteria according to its degree of relevance for management. The product of the weight and associated scores of the criteria returned the final score of the different abundance estimation methods. The final score was calculated so that the greater it was, the most suitable and relevant it was for duiker population management.

| Method | c1 | c2 | c3 | c4 | c5 | c6 | c7 | c8 | Score |
| --- | --- | --- | --- | --- | --- | --- | --- | --- | --- |
| Call screaming | 1 | 2 | 2 | 2 | 3 | 3 | 3 | 0 | 56.9 |
| Batting/Catching with nets | 1 | 3 | 1 | 3 | 2 | 2 | 2 | 2 | 54.2 |
| Nocturnal visual counting | 1 | 3 | 2 | 3 | 3 | 2 | 3 | 3 | 66.7 |
| Diurnal Visual counting | 1 | 3 | 2 | 3 | 2 | 2 | 2 | 3 | 58.3 |
| Counting dung | 1 | 1 | 1 | 1 | 2 | 2 | 2 | 0 | 34.7 |
| Indirect Recce | 1 | 1 | 1 | 1 | 2 | 3 | 3 | 2 | 44.4 |
| Camera trap | 1 | 3 | 2 | 1 | 1 | 2 | 2 | 0 | 41.6 |
| Tracks count | 1 | 1 | 0 | 0 | 3 | 2 | 3 | 0 | 36.1 |

**Key:** c1, the method have a good sampling design. We suppose that all methods are good sampling (c1 = 1); c2, the method identifies the species: if very difficult (c2=0), if difficult (c2 = 1), if good (c2 = 2) and if very good (c2 = 3); c3, the influence of the method on environmental factors: if very sensitive
(c3 = 0), if sensitive (c3 = 1), if less sensitive (c3 = 2) and if not sensitive (c3 = 3); c4, the risk of method counting twice the same species: if very high (c4 = 0), if high (c4 = 1), if lower (c4 = 2) and if almost zero (c4 = 3); c5, the technology used by method: if complex (c5 = 0), if difficult (c5 = 1), if easy (c5 = 2) and if very easy (c5 = 3); c6, the time used by the method to collect data and information, including analysis and processing: if very long (c6 = 0), if long (c6 = 1), if short (c6 = 2) and if fast (c6 = 3); c7, the material and financial resources used by the method: if very expensive (c7 = 0), if expensive (c7 = 1), if less expensive (c7 = 2) and if not expensive (c7 = 3); c8, the ability of the method to estimate population density or absolute abundance: if very good (c8 = 3), if good (c8 = 2), if less good (c8 = 1), and if almost zero or very bad (c8 = 0).

**FIGURE 2:**
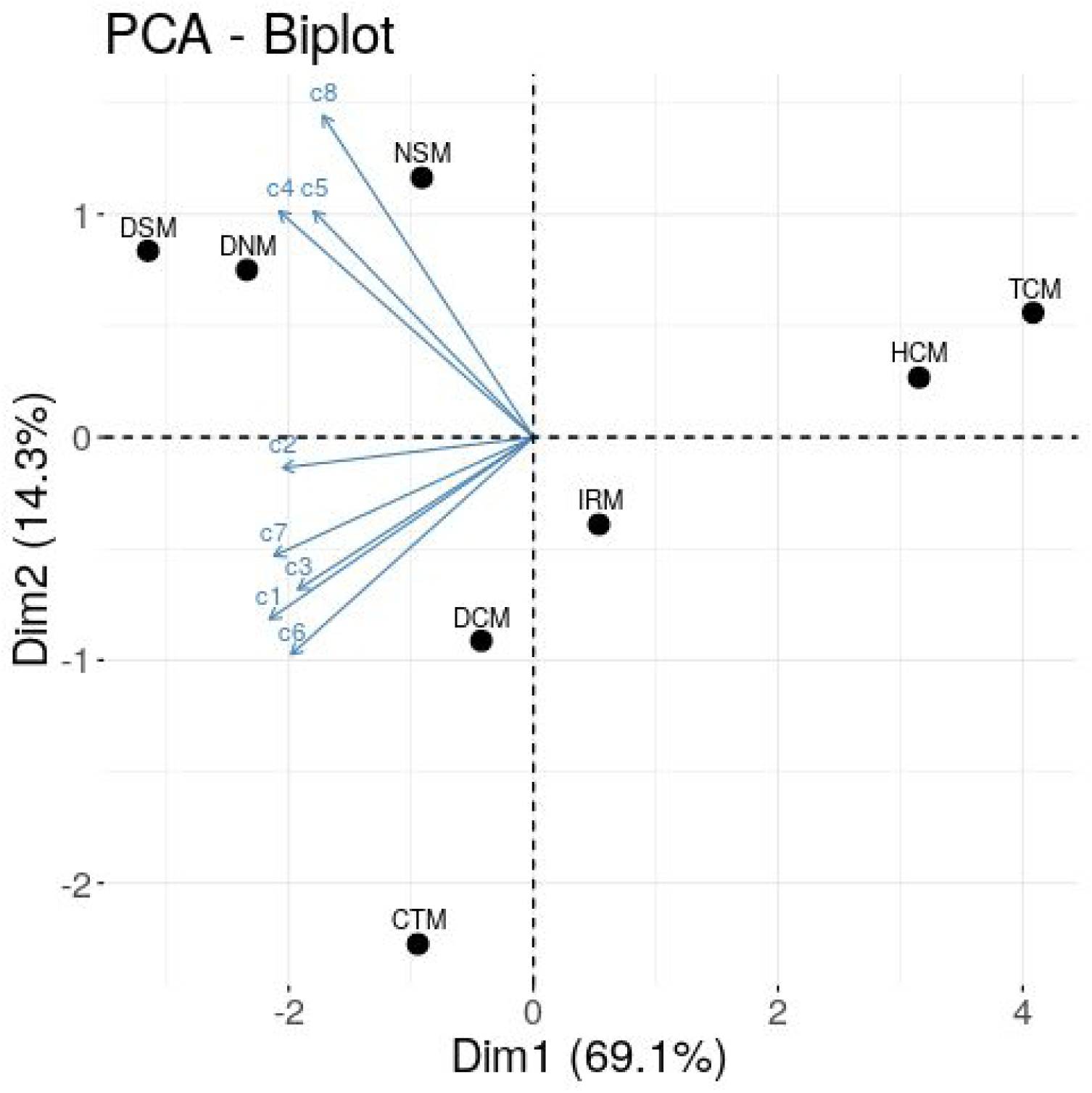
Principal component analysis (PCA) of methods for estimating abundance of duiker populations in African rainforest. Two principales components are highlighted: the first principal component (Comp. 1) is associated to criteria c8, c4 and c2 and opposed methods estimating absolute density (NSM, DSM, DNM), with simpler methods estimating relative abundance only (CTM and DCM). The second principal component (Comp. 2), associated to criteria c3, c5, c6 and c7, ranked methods on a gradient ranging from cheap and quick to carried out, to more labor consuming ones, and opposed estimating relative method (CSM and IRM). Hunter call method (HCM); Drive-netting method (BNM); Nocturnal sampling method (NSM); Diurnal sampling method (DSM); Dung count method (DCM), Indirect Recce (IRM).

## 4 DISCUSSION

The current absence of reference methods for estimating the abundance of mammals hunted for their bushmeat is at the heart of the questions of sustainable management in African rainforests (Bastin & Marechal, 2009; Fa & Brown, 2009; van Vliet et al., 2012; Wilkie & Carpenter, 1999; van Vliet & Nasi, 2008). Identifying a reliable method to monitor the abundance of duikers for management purposes is therefore crucial in the context of adaptive and sustainable management of wildlife and bushmeat. Ideally, such a method should be cheap, adapted to the environmental context while guaranteeing a sufficiently accurate estimate of the distribution and abundance of duiker species to capture temporal trends in abundance (Mathot & Doucet, 2006).

From our review and scoring of previously used methods, the distance sampling offers the best compromise in terms of costs and accuracy for estimating abundance of duikers, especially when sampling is carried out at night (NCM). Nocturnal observations are arguably the most efficient way to detect duikers because the spotlight reflects into the duikers’ eyes even if the visibility in dense rainforests is reduced and the animals are shy and stay hidden when alerted (Griffiths, 1979; Kamgaing et al., 2018; Lannoy et al., 2003; Payne, 1992). Notwithstanding its high relevance for management purposes, the distance sampling methods have been criticized because of “the difficulty of detecting all individuals on the transect midline” (Duckworth, 1998), which violates the main assumption of a perfect detection on the transect line (Buckland et al., 2004). However, thanks to further statistical developments, this assumption may be relaxed by, for instance, combining a standard distance sampling method with a double observer survey (Borchers, Laake, Southwell & Paxton, 2006; Kissling & Garton, 2006). Clearly, minimal alteration of the sampling design can dramatically improve the robustness of the distance sampling methods with little extra costs in terms of logistic and organization in the field, and in manpower as well, making distance sampling a reference for estimating the abundance of duiker populations.

Although accurate estimates of duiker abundance may be obtained using camera trap based (CTM) and CMR methods, these were not the most relevant for management purposes. Camera traps are an efficient tool to ensure continuous sampling and to work in areas difficult to access (Trolliet et al., 2014) and when individuals can be identified quite well (Foster & Harmsen, 2011; Trolliet et al., 2014), especially with upcoming artificial intelligence that will dramatically simplify and speed up the processing of pictures (Norouzzadeh et al., 2018). Even if the estimation of absolute abundance is theoretically possible using camera trap data, this option essentially applies to species with idiosyncrasies (*e.g.* Karanth et al., 2006 on tigers). Abundance estimates should be taken more cautiously for species for which individual recognition is not possible as risks of double counts remains high (Oliveira-Santos et al., 2009) and the estimation of detection rate is not readily available. Abundance estimation methods based on animal movement and ideal gas diffusion models have been proposed (e.g. spatially explicit capture-mark-recapture: Borchers & Efford 2008) but their performance remains to be assessed (Foster & Harmsen, 2012). Hence, methods based on camera traps currently lie in between absolute and relative abundance methods, such as the number of individuals seen per time units (Bowkett et al, 2007). Nevertheless, a complete setting of camera traps is still expensive and complicated to deploy in the field for wildlife managers (Bowkett et al., 2007; Carbone et al., 2001; van Vliet et al., 2009). Similarly, despite CMR methods are often regarded as a gold-standard in ecological studies, the repeated observation of the same individuals is very costly to implement and difficult to apply at the large spatial scale required for population management. Consequently, the drive-netting method did not score high for estimating the abundance of duikers (Table 3) and, if the non-invasive version of the CMR could be well adapted to these discrete nocturnal or low density species, one may face technical difficulties such as the lack of primers or the conservation of DNA in the tropical environment of Central Africa (Lukacs & Burnham, 2005; Mills et al., 2000; Petit & Valérie, 2006). Finally, even if the costs are continuously decreasing, DNA extraction and amplification on large sample sizes require substantial amount of money and cannot be proposed on the long run or for management agencies with often very limited funding capacities.

Among the relative abundance methods, we found the hunter call method (HCM) to be a relevant alternative to assess relative abundance of duiker populations. The main reason for why this method scores relatively good is its very simple implementation that does not require special skills if the observer is accompanied by an experienced hunter or use the adapted equipment (Newing et al., 2004). This method remains to be improved though, and particularly regarding its currently poor sampling design (Wilson, 1990). A well planned sampling design along linear transects with pre-defined timing of observation and repetition of transects are to be advised. Finally, because the method does not allow for imperfect detection, the absolute abundance of duikers remains unknown and the method is somewhat sensible to any changes in the conditions of observations such as habitat openness.

Conversely, the dung count method (DCM) received the lowest relevance score and hence, is considered as the least effective method for estimating the abundance of duiker populations inhabiting rainforests. This result is inconsistent with the statements of many authors (Newing et al., 2004; White & Edward, 2001) who concluded that dung counts offer a relevant method for estimating abundance of mammals, mainly because dungs are the most frequent and easier to find indice of animal presence. Yet, its low level of relevance is explained by the fact that dug counts present major methodological pitfalls (Marechal, 2011; Mathot & Doucet, 2006; Plumptre, 2000; van Vliet & Nasi, 2008; van Vliet et al., 2009; White & Edwards, 2001). For instance duikers mark their territory with feces so that dung distribution is not random, and the rate of defecation and of decomposition of the dung are two variables that are difficult to estimate, which prevents a reliable estimate of the abundance of duiker populations (Plumptre, 1995; Nasi & van Vliet, 2011). We acknowledge that these limitations should not hamper the relevance of sampling feces to assess the presence of mammalian species in a given area.

Regardless the technical issues associated with each method we reviewed, getting estimates of absolute or relative abundance of duikers do not provide information about the demographic processes that are important for population management such as density-dependence. Density-dependence is notoriously known to be difficult to detect from count data (Dennis & Tapper, 1994; Knape & de Valpine, 2011). Changes in population dynamics are likely easier to be revealed by a reduction in reproductive rate of females or a decrease in the body mass of juveniles (see Bonenfant et al., 2009 for a review). To overcome the technical problems and interpretation limitations emerging with any abundance estimate, we propose to combine estimates of relative abundance with additional biological measurements that can be collected from harvested duikers such as body mass, as done within the framework of the indicators of ecological changes (IEC, Morellet et al., 2007). This approach first developed for managing roe deer (*Capreolus capreolus*) populations (Cederlund et al., 1998) has been successfully deployed on other species of large herbivores in the holarctic region (see Garel et al., 2010 for an example on red deer *Cervus elaphus*). The basic principle of IEC is built on the concept of density-dependence: at a certain level of density, the resources available to an individual diminish and can lead to biological changes in its survival, reproduction, physical performance which ultimately affects abundance (Bonenfant et al., 2009). ICE have been designed to capture these changes with easy-to-measure proxies (Morellet et al., 2007). Currently, three families of IEC are distinguished: (*i*) ICE providing information on the relative abundance of the target population; (*ii*) IEC reflecting changes in the phenotypic quality of individuals; and (*iii*) IEC providing information on the pressure of individuals on vegetation. A joined monitoring of ICE including all these three families is required to manage the ungulate-environment system. This framework fits the principle of adaptive management based on a trial-and-error process. By applying IECs over time managers will progressively adjust the hunting bags to the changes in IEC revealing the functioning of the ungulate-environment system. As information accumulates, it becomes possible to refine the hunting quotas.

In addition to the distance sampling or indirect sampling methods by single observation on individuals based the indices to monitor population abundance, three IECs could be proposed for the management of duikers in the context of tropical rainforests of Central Africa: body mass, jaw length and a fruit consumption index. The first two IECs describe the physical condition of duikers, which reflects variations in individual performance with lighter duikers having a lower per capita food intake (*i.e.* too many animals or too scarce food resources). Although duikers are not officially hunted, the monitoring of IEC indicating performance could be applied in tropical forests. One option would be a regular sampling and measurements of hunted duikers sold at different local markets. Indeed, such markets are often located near protected areas, village hunting areas or communal hunting grounds. Similarly to the consumption of shrubs by deer, the consumption of forest fruits and seeds (*e.g. Irvingia gabonensis, Pseudospondias longifolia, Dacryodes buettneri, Dacryodes edulis, Ancystrophyllum spp.* (Gautier-Hion, Emmons & Dubost, 1980; Feer, 1989, 1995) by duikers could be measured by quantifying the amount of such food on the ground across years. However, in the African rainforest conditions, many species of large herbivores are sympatric and the plant phenology (fructification, flowering, leafing, etc.) is complex, likely making the IEC indicating the impact on vegetation difficult to interpret and less relevant than those indicating population abundance and individual performance.

## 5 CONCLUSION

We found that four out of seven methods previously applied to duiker populations may be recommended to estimate duiker abundance in tropical forests. However, in spite of their interest for management purposes, these methods are still limited in the context of large-scale population management because they do not provide information about population dynamics (*e.g.* density-dependence processes). For a more meaningful monitoring of duikers, we propose to cast these population abundance methods within the framework of indicators of ecological change (Morellet et al., 2007). A longitudinal monitoring of IECs and clear management goals (e.g. increase vs. control of wildlife populations) could help meeting the potential needs of rural and urban populations relying of bushmeat on the one hand, and the needs to manage resources while respecting environment on the other hand. We contend that the IEC approach remains to be implemented in the different duiker hunting areas in Central Africa. Thus, once this association is validated scientifically, the Central African wildlife management institutions of each country, but also the Central African Forest Commission (COMIFAC) which is a sub-regional institution of concerted management policy at the level of ten Central African countries, could promote the management of duikers based on IEC, its implementation in the field, as well as the analysis and interpretation of the data.

### ORCID

*Christophe Bonenfant http://orcid.org/0000-0002-9924-419X Jean-Michel Gaillard https://orcid.org/0000-0003-0174-8451*

## ACKNOWLEDGEMENTS

We thank the Government of the Republic of France through the Campus France program, the Government of the Democratic Republic of Congo (DRC) through the Ministry of Environment and Sustainable Development and The Ministry of Higher Education and University, the Program for the Conservation of Forest Ecosystems in Central Africa (PACEBCo) of the Central African Forests Commission (COMIFAC) and the Nature Plus of Belgium, for their financial, technical, logistical and didactic support. Our thanks also go to Punga Kumanenge Julien and Palata Jean-Claude for their collaboration. Finally, we thank Cayla Fabien from the Embassy of France in the Democratic Republic of Congo- Kinshasa for its support and help.

